# Allele Dispersion Score: Quantifying the range of allele frequencies across populations, based on UMAP

**DOI:** 10.1101/2022.02.11.479970

**Authors:** Solenne Correard, Laura Arbour, Wyeth W. Wasserman

**Affiliations:** BC Children’s Hospital Research Institute, University of British Columbia, Vancouver BC, Canada; Centre for Molecular Medicine and Therapeutics, Department of Medical Genetics, University British Columbia, Vancouver BC, Canada; Department of Medical Genetics, University of British Columbia, Vancouver BC, Canada; Division of Biomedical Sciences, University of Victoria, Victoria, British Columbia, Canada

## Abstract

Genomic variation plays a crucial role in biology, serving as a base for evolution - allowing for adaptation on a species or population level. At the individual level, however, specific alleles can be implicated in diseases. To interpret genetic variants identified in an individual potentially affected with a rare genetic disease, it is fundamental to know the population frequency of each allele, ideally in an ancestry matched cohort. Equity in human genomics remains a challenge for the field, and there are not yet cohorts representing most populations. Currently, when ancestry matched cohorts are not available, pooled variant libraries are used, such as gnomAD, the Human Genome Diversity Project (HGDP) or the 1,000 Genomes Project (now known as IGSR: International Genome Sample Resource). When working with a pooled collection of variant frequencies, one of the challenges is to determine efficiently if a variant is broadly spread across populations or appears selectively in one or more populations. While this can be accomplished by reviewing tables of population frequencies, it can be advantageous to have a single score that summarizes the observed dispersion. This score would not require classifying individuals into populations, which can be complicated if it is a homogenous population, or can leave individuals excluded from all the predefined population groups. Moreover, a score would not display fine-scaled population information, which could have privacy implications and consequently be inappropriate to release. Therefore, we sought to develop a scoring method based on a Uniform Manifold Approximation and Projection (UMAP) where, for each allele, the score can range from 0 (the variant is limited to a subset of close individuals within the whole cohort) to 1 (the variant is spread among the individuals represented in the cohort). We call this score the Allele Dispersion Score (ADS). The scoring system was implemented on the IGSR dataset, and compared to the current method consisting in displaying variant frequencies for several populations in a table. The ADS correlates with the population frequencies, without requiring grouping of individuals.

## Introduction

Genomic variation plays a crucial role in biology, serving as the basis for adaptation in populations and evolution of species. While most variants are benign, on the individual level, some DNA variants can modify risk or cause disease. For clinical interpretation of variants, it is crucial to gather as much relevant information as possible when studying a variant, such as the functional role(s) of the location at which it appears (i.e. a coding gene, a regulatory region, etc) (Zerbino et al., 2020), the conservation of the allele across several species (Lindblad-Toh et al., 2011), and the frequency of the given allele in a reference cohort (Oak et al., 2020; Whiffin et al., 2017). In every species with an available reference genome, as the number of sequenced genomes grows, allele frequencies become more reliable and population frequencies emerge.

In human, the largest collection of allele variation data is gnomAD, coalescing genomic data from over 75,000 individual whole genomes in gnomAD v3 (Karczewski et al., 2020). Even as one of the largest pools of genomes, some populations lack representation. In order to improve representation of populations, sequencing a sufficient number of individuals from ancestrally diverse backgrounds, in ethical and community-based initiatives is necessary. Initiatives are underway to pursue balanced representation (Caron et al., 2020; Fattahi et al., 2019; Lee et al., 2017). As representation of populations increases, it will become increasingly common to consider if an observed variant is spread across one or a few population(s) or broadly across multiple populations. The capacity to efficiently describe such dispersion of alleles across populations is the focus here. In this report, populations refer to groups of individuals that are genetically close, which are sometimes referred to as sub-population in other reports.

Currently, a widely used method to indicate if a variant is narrowly or broadly spread across populations, is to create population categories, as in gnomAD where ten populations are identified. This method requires assigning each individual to a population. Several methods exist to assign individuals to a population as it can become challenging as cohort sizes grow (Gimbernat-Mayol et al., 2021). gnomAD used a random forest classifier trained using 16 principal components (PCs) as features on samples with known ancestry. Ancestry was then assigned to all samples for which the probability of that ancestry was > 75% according to the random forests model, and all remaining samples were assigned as “other” (oth). One of the main advantages of this model is the capacity to report a table containing a variant’s frequency for each of the constituent populations. However, it is only possible when some samples are of known ancestry to train the model, and it leads to the classification of a subset of outlying samples as “other” (5% of samples in gnomAD overall v.3 [1,047 / 20,744]).

Another challenge is that fine-scale population mapping data may not be acceptable to release because of privacy and concerns with data sovereignty (Carroll et al., 2021; Hudson et al., 2020). Following problems of discrimination faced by indigenous peoples throughout the world, including oppression, marginalization and research abuses, the United Nations recognized the right for Indigenous people to control the use of their data (UNDRIP, 2007). Hence, separating the individuals present in a genomic database into populations and sharing population specific information could be an issue for some Indigenous populations, and therefore, a method addressing this concern was sought. The resulting method consists in associating a single score with each variant that indicates if the variant is spread broadly or focused locally without revealing specific distribution data or population information.

To address the challenge of succinctly describing the dispersion of an allele across populations, we sought to develop a scoring method where, for each allele, the score can range from 0 (the variant is limited to a subset of close individuals within the whole cohort) to 1 (the variant is spread among the individuals present in the cohort). This score, called the Allele Dispersion Score (ADS), is associated with each allele, complementing the overall allele frequency. As for the allele frequency, the ADS will differ depending on the cohort studied. The ADS is calculated based on projected distance between individuals carrying the variant within a Uniform Manifold Approximation and Projection (UMAP). The ADS score was developed and tested on the International Genome Sample Resource (IGSR) dataset, and correlates with the population’s frequencies. It proved to be a metric capturing the allele dispersion for biologically relevant variants, without displaying population specific information or requiring grouping of individuals. We believe that the ADS should be displayed together with the Minor Allele Frequency (MAF) when developing a background variant library or reporting the characteristics of a variant.

## Results

The calculation of the ADS (fully detailed in the Materials and Methods section) is a two steps process: (i) a UMAP is created based on a random subset of genotype data from all individuals. In the projection, individuals that are genetically similar will be grouped (Diaz-Papkovich et al., 2019, 2021); (ii) for each variant, the ADS is calculated based on the projected distance between the individuals carrying the alternate allele. Hence, if the allele is limited to a population (i.e. individuals that are close genetically), the distance between the individuals will be small, and the ADS will be on the lower end of the range. Even though UMAP is a non-linear representation of distance, its capacity to represent individual proximity is particularly well-suited to this problem.

### UMAP generation

The ADS, which represents the spread of a given allele across individuals within a dataset, is calculated based on a UMAP. UMAPs have previously been used in population genomics to study population structure (Diaz-Papkovich et al., 2021). Using the IGSR dataset (Clarke et al., 2017) and PLINK (Chang et al., 2015), the 15 first principal components were calculated and then a UMAP was generated using the R UMAP package (McInnes et al., 2020). Individuals can be colored according to their continental label (Figure 1A) or population label (Figure 1B).

**Figure 1.**
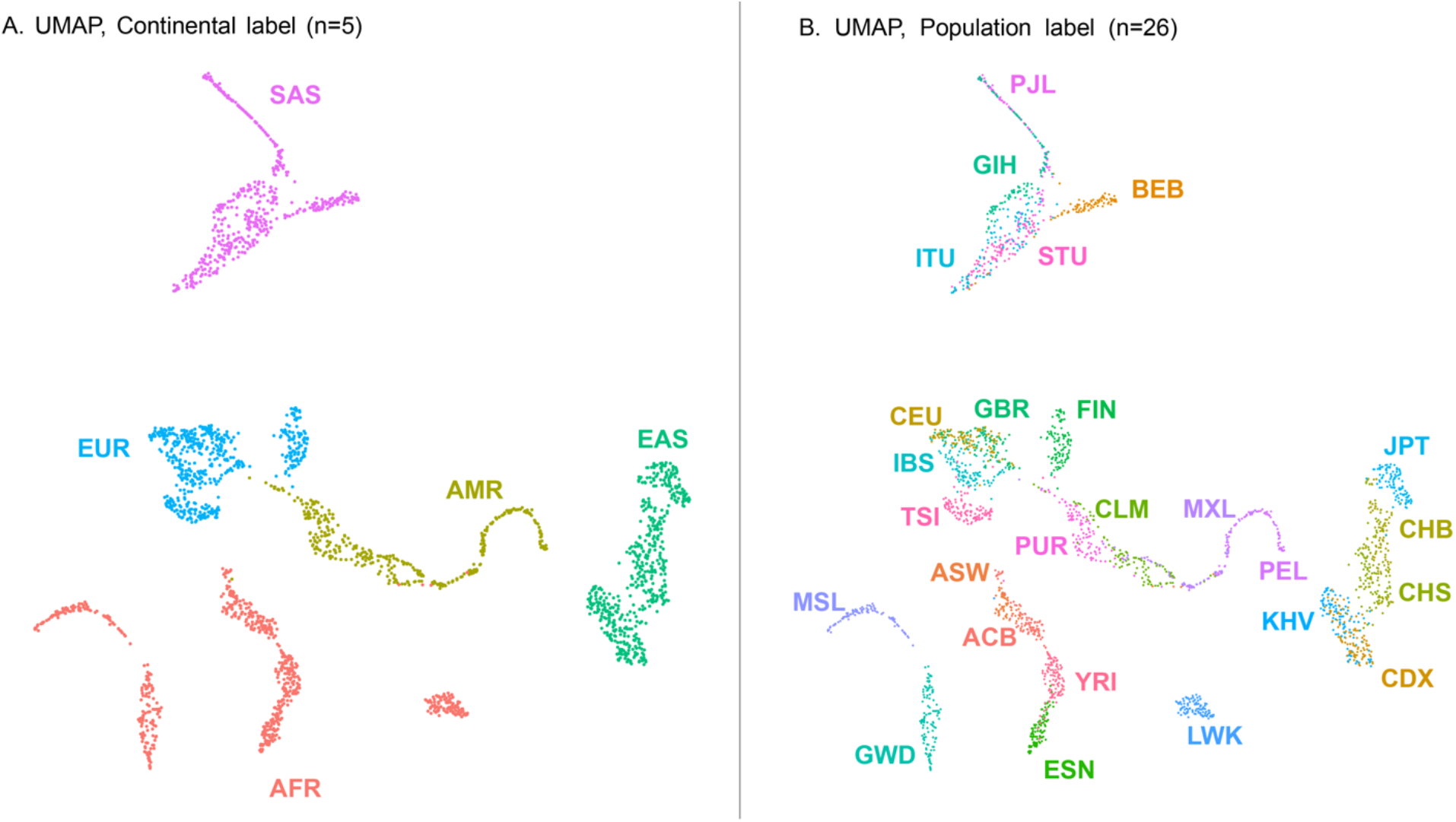
UMAP obtained with the first 15 principal components of the IGSR data. (A) Individuals colored according to their continental label as assigned in the IGSR dataset. AFR, African; AMR, Ad Mixed American; EAS, East Asian; EUR, European; SAS, South Asian. (B) Individuals colored according to their population label as assigned in the IGSR dataset. ACB, African Caribbean in Barbados; ASW, African Ancestry in Southwest US; BEB, Bengali; CDX, Chinese Dai; CEU, Utah residents with Northern/Western European ancestry; CHB, Han Chinese; CHS, Southern Han Chinese; CLM, Colombian in Medellin, Colombia; ESN, Esan in Nigeria; FIN, Finnish; GBR, British in England and Scotland; GWD, Gambian; GTH, Gujarati; IBS, Iberian in Spain; ITU, Indian Telugu in the UK; JPT, Japanese; KHV, Kinh in Vietnam; LWK, Luhya in Kenya; MSL, Mende in Sierra Leone; MXL, Mexican in Los Angeles, California; PEL, Peruvian; PJL, Punjabi in Lahore, Pakistan; PUR, Puerto Rican; STU, Sri Lankan Tamil in the UK; TSI, Toscani in Italy; YRI, Yoruba in Nigeria. Axes in UMAP are arbitrary.

### Allele Dispersion Score and other population segregation

The ADS is calculated for each variant with an allele count (AC) of at least two in the dataset. The ADS, which ranges from 0 (not spread) to 1 (maximally spread), is calculated based on the relative distance between the individuals carrying the allele of interest in the UMAP. As individuals more genetically similar should be proximal on the UMAP, if a variant is limited to a population, the relative distance between the individuals with the allele will be small, compared to an allele present in the same number of individuals, but not restricted to a population.

Using the IGSR dataset, the ADS was calculated for a total of 45,183,262 variants. Even though the ADS could in theory vary from 0 to 1, a non-homogeneous distribution was expected as real data were used. Using the IGSR dataset, the observed ADS vary from 0 to 0.65 (Figure 2A). The ADS, even though normalized with the theoretical lowest and highest values for the same allele count (AC), is associated with the variant allele frequency (Fig 2B and 2C). Within the IGSR dataset, as expected, variants with a low minor allele frequency (MAF) tend to have a lower ADS and variants with a higher frequency tend to have a higher ADS. To better interpret if a given variant is spread across a population using the ADS, the variant’s ADS can be compared to the ADS distribution for variants with a comparable allele frequency within the same dataset. The values of the ADS do not correlate with the genomic annotation of the variant (Figure 2D).

**Figure 2.**
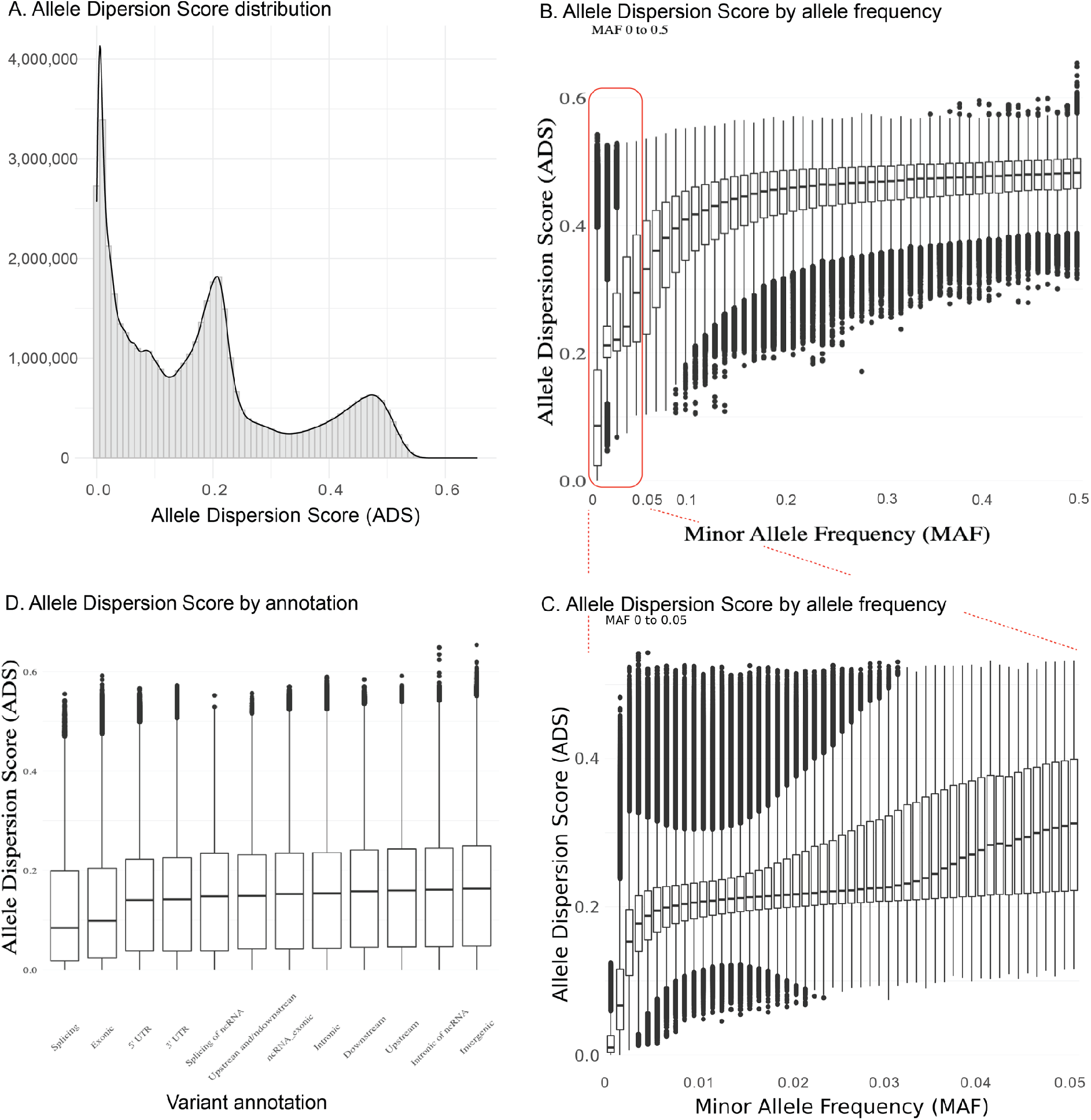
Distribution of the Allele Dispersion Score and distribution of the Allele Dispersion Score within bins grouping variants according to their MAF or their annotation for 45,183,262 variants from the IGSR dataset. (A) Histogram representing the ADS distribution. The ADS could vary from 0 to 1. Within this dataset, it varies between 0 and 0.65. (B) ADS distribution according to the Minor Allele Frequency (MAF). Variants are binned according to their MAF (bins of 0.01), the number of variants per bin is available in Table A in S2 Tables. The ADS correlates with the MAF. The first five bins (MAF < 0.05) circled in red are presented in Figure C (C) Zoom in a subset of bins (MAF < 0.05, bins of 0.001), the number of variants per bin is available in Table B in S2 Tables. (D) ADS distribution according to the variant annotation in the genome. Only annotations with more than 500 variants are represented. The number of variants per annotation is presented in S3 Table and the distribution per MAF for each annotation is presented in S4 Figure.

### Comparing the Allele Dispersion Scores with the number of population observations

A current approach for assessing a variant’s dispersion is to review the variant’s MAF in each population. Within the IGSR data, each individual is associated with two population labels: i) a continental label or super-population (n = 5); and ii) a population label (n = 26). The number of individuals per population is not evenly distributed (S5 Tables). Based on these groups of individuals, we calculated a label score, which is, for each variant, the number of populations with at least one individual carrying the alternate allele (can vary from 1 to 5 for the continental label and from 1 to 26 for the population label). Interestingly, with the continental label, 41.7% of the variants (18,871,497/45,183,262) have a label score of 1, meaning that they are present in only one population (Figure 3A, lower histogram) while, with the population label, most variants (18%, 8,140,806/45,183,262) have a label score of 2 (Figure 3C, lower histogram). This is due to the fact that many populations are not distinguishable, such as KHV and CDX, or ASW and ACB, as visible on the UMAP (Figure 1B).

**Figure 3.**
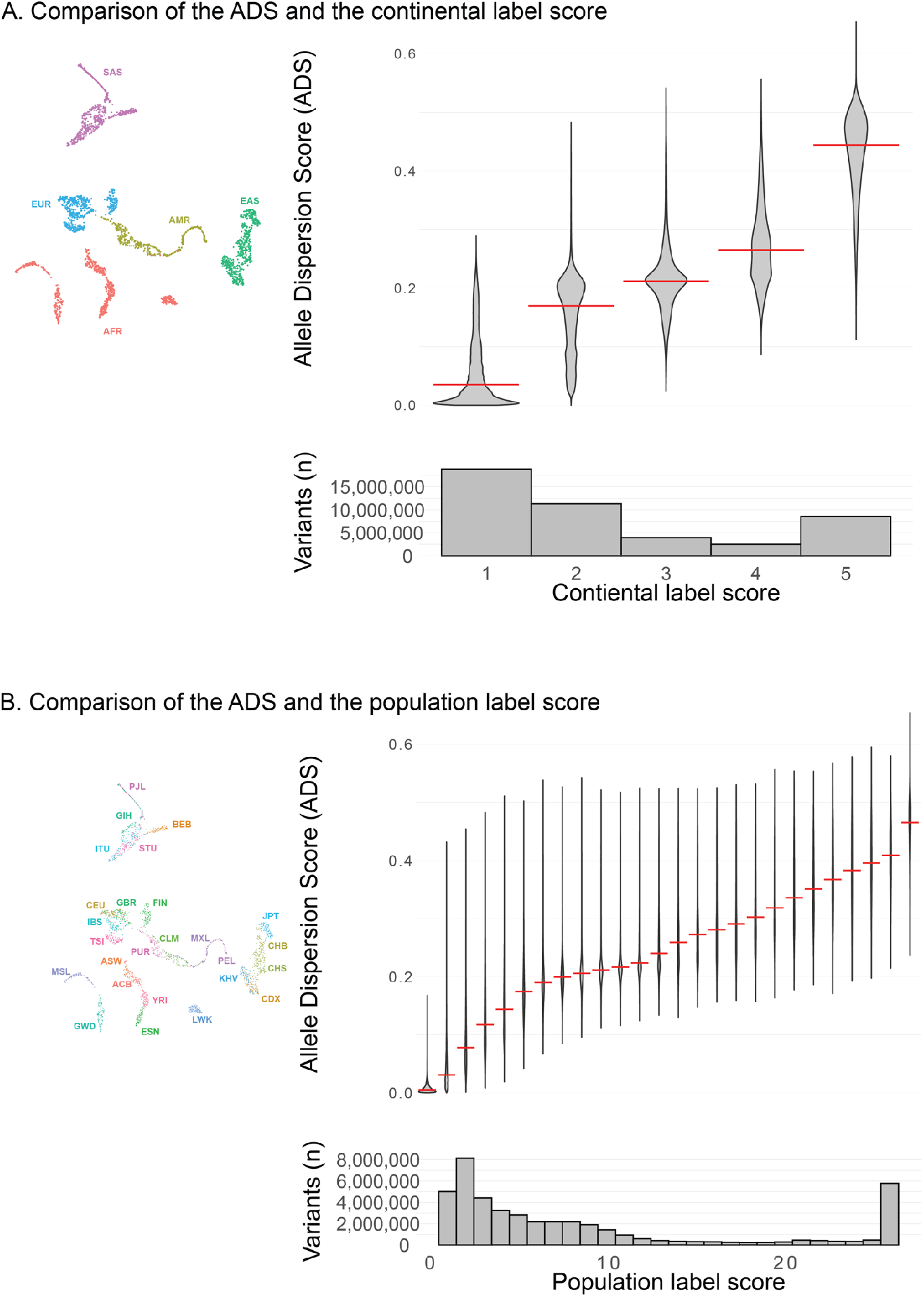
The Allele Dispersion Score correlates with the label score. (A) Upper left : UMAP used to calculate the ADS. The individuals are colored according to their continental label. Upper right : Violin plots showing the ADS (y-axis) compared to the continental label score (x-axis). The label score corresponds to the number of populations with at least one individual carrying the allele of interest. The red line represents the median ADS value. Lower panel : Distribution of the continental label score. The maximal number of populations is reached (n = 5). (B) Upper left : UMAP used to calculate the ADS. The individuals are colored according to their population label. Upper right : Violin plots showing the ADS (y-axis) compared to the population label score (x-axis). The red line represents the median ADS value. Lower panel : Distribution of the population label score. The maximal number of populations is reached (n = 26).

A Pearson product-moment correlation coefficient was computed to assess the relationship between the ADS and the label score. For both the continental and the population label score, there is a positive correlation and the relation is significant (Continental label and median ADS : Figure 3A upper right panel, Pearson’s correlation, r = 0.97, p-value = 0.007, Population label and median ADS : Figure 3B, upper right panel, Pearson’s correlation, r = 0.98, p-value < 2.2e-16). The full results of the statistical analysis are presented in S6 Figure.

### Representation of four extreme Allele Dispersion Scores

In order to better convey what is the actual allele dispersion represented by the ADS, four variants are shown in figure 4. To compare appropriately, variants with very different ADS but similar frequencies were selected. Each panel represents the UMAP obtained with the IGSR dataset, where each of the 2,548 individuals are colored according to their genotypes. As the ADS correlates with the variant frequency, each panel includes the ADS distribution for variants with a MAF in the same range as the MAF of the variant of interest (MAF +/− 10%). The ADS of the variant of interest is represented by a red dotted line. Panels A and B represent the distribution of two different variants within the dataset, with a comparable elevated MAF (frequent variant, MAF = 0.26) but different ADS : 0.21 (Figure 4A) and 0.56 (Figure 4B). Panels C and D represent two different variants, with, this time, a comparable low MAF (rare variant, MAF = 0.0011) but different ADS : 0.0089 (Figure 4C) and 0.48 (Figure 4D) (Table 1).

**Figure 4.**
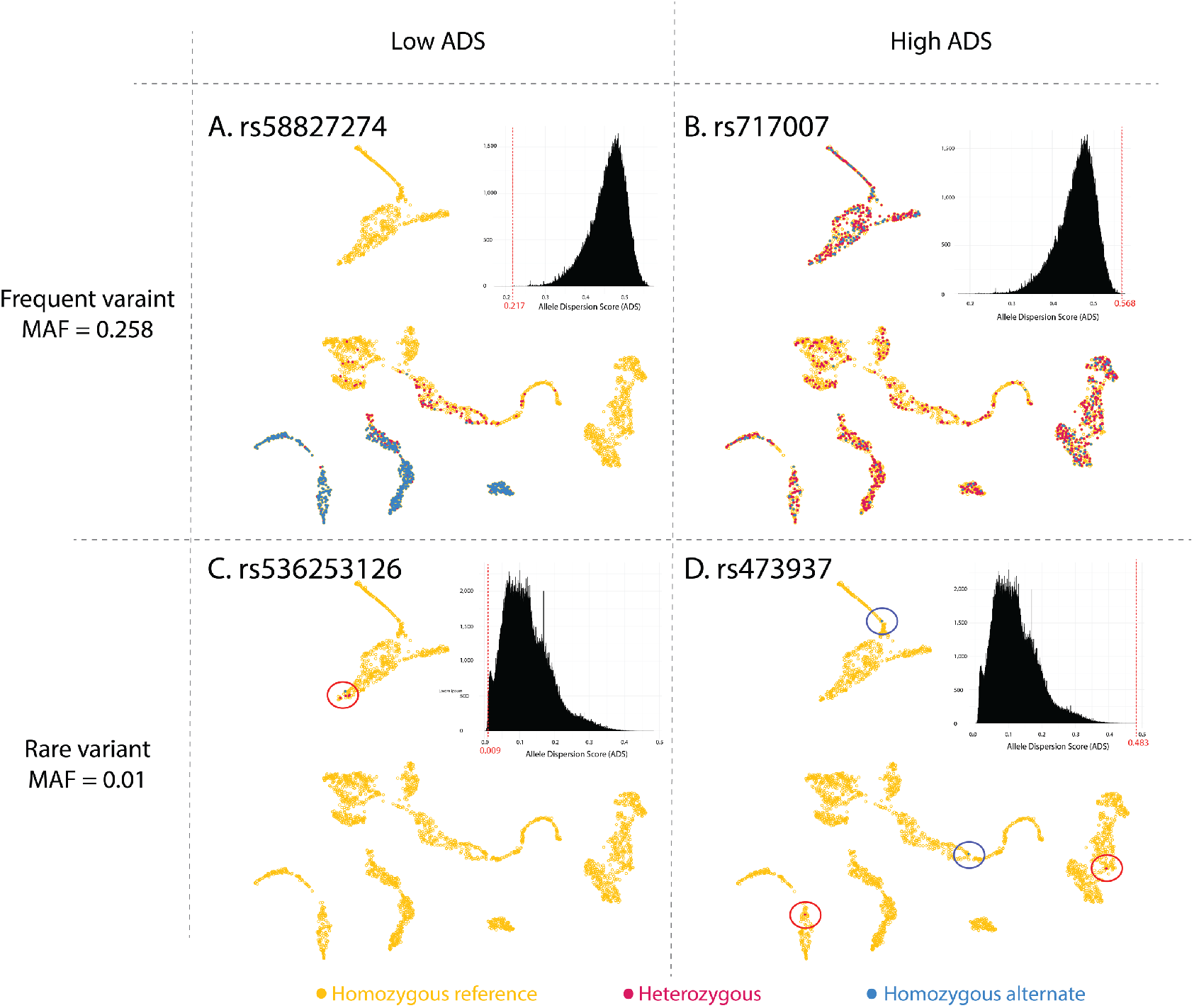
Visualization of the UMAP and the ADS for four specific variants. (A)-(D) Case studies of individual variants (described in Table 1) showing the dispersion of the allele within UMAP plots and inserting the variant ADS within ADS distributions. Panels A and B represent two variants with a high MAF (0.26), while panels C and D represent variants with a low MAF (0.0011). Each panel represents the UMAP obtained with the IGSR data where each individual is colored according to its genotype for a specific variant. Yellow : Individuals homozygous for the reference allele, Blue : Individuals homozygous for the alternate allele, Red : Individuals with heterozygous alleles. In panels C and D, the individuals carrying the variant of interest (heterozygous or homozygous alternate) are circled. Upper left corner of each UMAP: ADS distribution for variants with a comparable MAF than the variant of interest. The red score and dotted line represent the ADS of the variant of interest.

**Table 1.** Details of the variants presented in Figure 4. Chr : Chromosome, Ref : Reference allele, Alt : Alternate allele, MAF : Minor Allele Frequency. The gnomAD label score is obtained by counting the number of populations with at least one individual carrying the allele of interest.

| Panel<br>(Fig. 4) | chr | position<br>(GrCh 38) | Ref | Alt | dbSNP ID | MAF |  | Scores |  |  |  |
| --- | --- | --- | --- | --- | --- | --- | --- | --- | --- | --- | --- |
|  |  |  |  |  |  | 1KG<br>dataset | gnomAD<br>(v3) | ADS | Population<br>label score<br>(max = 26) | Continental<br>label score<br>(max = 5) | gnomAD<br>label score<br>(v3, max=10) |
| A | 4 | 3664767 | C | G | rs58827274 | 0.258 | 0.232 | 0.217 | 15 | 3 | 9 |
| B | 7 | 132265538 | C | A | rs717007 | 0.258 | 0.151 | 0.568 | 26 | 5 | 10 |
| C | 6 | 116095673 | T | C | rs536253126 | 0.001 | 0.000 | 0.009 | 1 | 1 | 1 |
| D | 11 | 117138294 | A | G | rs473937 | 0.001 | 0.000 | 0.483 | 4 | 4 | 5 |

### Representation of four variants of biological relevance

As a proof of concept, the ADS was analyzed for four variants with various biological relevance. Variants implicated in rare diseases are unexpected to be present in the IGSR dataset, therefore, examples focus on variants associated to a specific phenotype, polygenic risks scores (PRS), or cancer predisposition. Indeed, when a variant is identified in a population as associated with a given phenotype, the ADS can indicate quickly if the variant is spread across genetically close individuals or widely across several populations.

- The first variant (Figure 5A), rs73885319 (GrCh38, chr22: 36265860 A>G), in *APOL1,* is associated with end-stage kidney disease (ESKD) and is known to be more frequent in the African / African-American populations (Lin et al., 2019). In gnomAD, this variant has an allele frequency of 0.06, however, it is more frequent in the African / African-American group (AF of 0.22) than in other populations (0.034 for the “other” group and lower for the other groups). Within the IGSR dataset, it has an allele frequency of 0.07 and an ADS of 0.14 (Table 2). Such an ADS is on the lower end of the ADS distribution for variants with comparable allele frequencies. The dispersion of the individuals carrying the C allele in the UMAP shows that, indeed, the variant is carried mostly by individuals labeled as AFR by the IGSR project (Figure 5, panel A).

**Figure 5.**
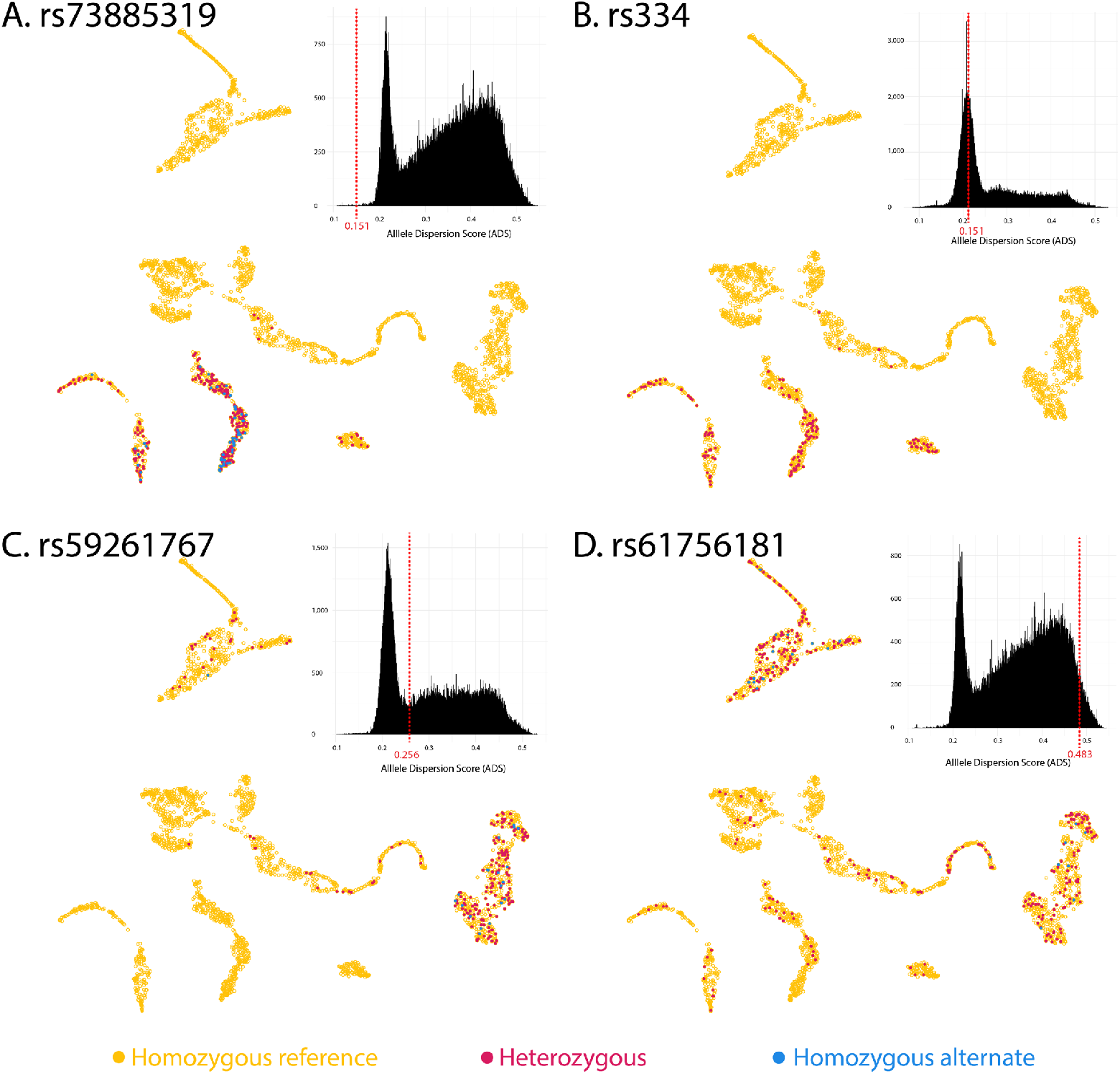
Visualization of the UMAP and the ADS for four biologically relevant variants. (A)-(D) Case studies of individual variants (described in Table 2) showing the dispersion of the allele within UMAP plots and inserting the variant ADS within ADS distributions. Each panel represents the UMAP obtained with the IGSR data where each individual is colored according to its genotype for a specific variant. Yellow : Individuals homozygous for the reference allele, Blue : Individuals homozygous for the alternate allele, Red : Individuals heterozygotes. In panels C and D, the individuals carrying the variant of interest (heterozygous or homozygous alternate) are circled. Upper left corner of each UMAP: ADS distribution for variants with a comparable MAF than the variant of interest. The red score and dotted line represent the ADS of the variant of interest.

**Table 2.**
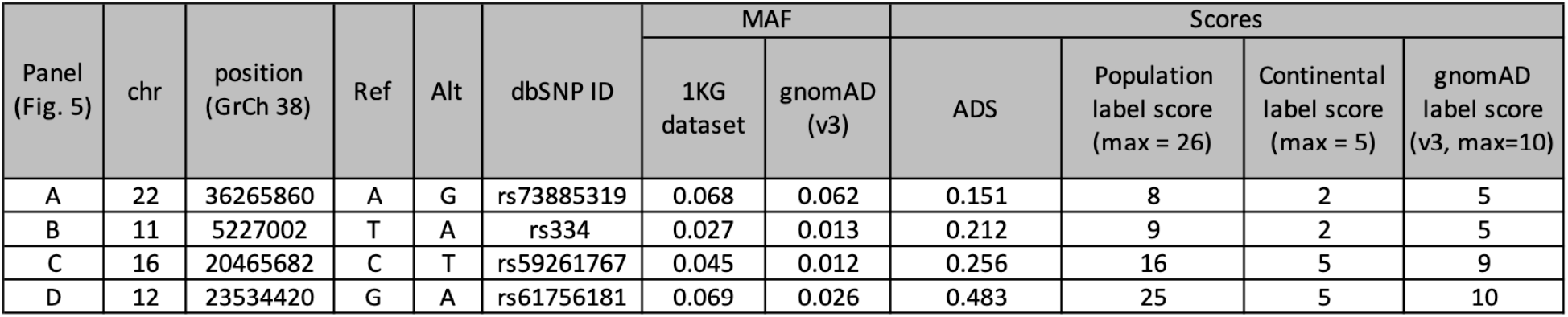
Details of the variants presented in Figure 5. Chr : Chromosome, Ref : Reference allele, Alt : Alternate allele, MAF : Minor Allele Frequency. The gnomAD label score is obtained by counting the number of populations with at least one individual carrying the allele of interest.

- The second variant, rs334 (GrCh38, chr11: 5227002 T>A) is a missense variant located in *HBB.* This variant is involved in Sickle cell disease, an autosomal recessive disease relevant to global health (Piel et al., 2017). Because of the global health implication of this variant, its origin as well as its frequency across different populations is under scrutiny (Kakande et al., 2020; Shriner and Rotimi, 2018). The global frequency of the variant is 0.012 in gnomAD, with a higher prevalence in the African / African-American population (MAF = 0.04) compared to other groups (MAF < 0.007). In the IGSR dataset, the global MAF is 0.027, and the ADS is 0.16, which is close to the peak in the distribution. In the UMAP, individuals carrying the alternate allele are mostly labelled as AFR. There are no homozygous individuals for this variant in the IGSR dataset.

- The third variant, a stop-gain in *ACSM2A* (rs59261767, GrCh38, Chr16:20465682 C>T) increases the concentration of indolepropionic acid in plasma and completely abolishes the catalytic activity of ACSM2A (Koshiba et al., 2020). This variant, and the metabolic phenotype associated, are known to be more frequent in the Japanese population. In gnomAD, this variant has a MAF of 0.012, while the MAF in the IGSR dataset is 0.045. This difference can be explained by the much lower proportion of East Asian individuals in gnomAD compared to the IGSR dataset (2,582/76,067 : 3.4% in gnomAD vs 515/2,548 : 20% in the IGSR dataset at this position).

The population specific distribution is clearly observed within gnomAD, as this variant has a higher MAF in the East Asian group (AF = 0.19) compared to other populations (AF < 0.037). The ADS calculated based on the IGSR dataset for this variant is 0.31. Based on the distribution of this allele on the UMAP, it is visible that the variant is more common in the EAS group (even if the JPT group does not seems to display more carriers than the other EAS groups), while some carriers are also present in other populations (Figure 5C).

- The fourth variant, rs61756181 (GrCh38, chr12:23534420 A>G), is a synonymous variant in *SOX5. SOX5* encodes a transcription factor implicated in Lamb-Shaffer syndrome, an autosomal dominant disorder (Nesbitt et al., 2015). Even though *SOX5* is implicated in the disease, rs61756181 is a relatively frequent variant in this gene (MAF of 0.026 and 0.069 in gnomAD and the IGSR dataset respectively), not known to be associated to any phenotype. This variant is present in all of the populations represented in gnomAD, with varying MAF (from 0.129 for the South Asian population to 0.001 in the Amish population). This presence across multiple populations is also visible through the ADS score of 0.496, which is on the high end of the distribution. When looking at the variant distribution on the UMAP, the presence of the variant in every group is also visible.

## Discussion

In clinical genetics, knowing the allele frequency and the number of homozygotes for a given allele across several populations is important. Information representing the spread of a given variant across populations is also of interest, as represented by several databases which separate their cohorts into populations. The Allele Dispersion Score (ADS) was created to assess the dispersion of a variant within individuals without the need to classify individuals into populations. This score calculation is based on a UMAP and was tested on the IGSR dataset.

### UMAP generation and visualization

A UMAP was generated using the 15 first principal components calculated based on 5% randomly selected variants on each chromosome of the IGSR final (phase 3) 2015 release including 2,548 unrelated samples. Each sample is associated with a continental and a population label, based on individual self-identification. Previously, Diaz-Papkovich et al. (2019), also published a UMAP using the IGSR dataset, however, they used genotype data from 3,450 individuals generated by Affy 6.0 SNP-chip genotyping. As we used a slightly different dataset, it was interesting to compare the two UMAPs.

Both UMAPs concur with some populations overlapping, such as CEU and GBR or KHV and CDX while some populations are distinct within both UMAPs such as LWK or BEB. However, some differences are visible, which can be explained by two non-exclusive reasons. First, the original dataset is slightly different for each, indeed, the Affy 6.0 SNP-chip was used by Diaz-Papkovich et al., while we used a subset of the variants obtained from whole genome sequencing. In our dataset the variants were selected randomly, and some may be less informative (i.e. less polymorphic) than the ones included in the SNP-chip. We believed the data randomly obtained from whole genome sequencing held the greatest relevance as it is more likely to be used in future projects generating the ADS. Secondly, we used a different software to generate the UMAP compared to the software used by Diaz-Papkovich et al., even though some parameters are similar, it may explain some of the variability.

### ADS interpretation

The ADS proves to be an efficient score to summarize the spread of a variant across populations. The label score, which is, for each variant, the number of populations with at least one individual carrying the alternate allele according to the IGSR population groups, correlates with the ADS (Figure 3). Moreover, using four biologically relevant variants, the ADS score captures the spread of the variants across the populations as expected (Figure 5). Finally, when identifying the individuals carrying the allele of interest on the UMAP together with the ADS, the ADS varies in a manner that is consistent with expectation.

### The ADS do not require grouping individuals into populations

Interestingly, the ADS calculation does not require any assignment of individuals into populations. In other methods, classifying can be done based on genomic data only (like gnomAD) or based on self-reporting (like for the IGSR labels). However, classifying individuals into populations has limitations when individuals are not associated with one specific population. As the number of individuals included in background variant libraries increases, it will be increasingly complex to group individuals into distinct populations, and this may lead to an increased number of individuals classified in “other”. Indeed, according to gnomAD, “Individuals were classified as ‘other’ if they did not unambiguously cluster with the major populations in a principal component analysis (PCA)”, which currently represents 5% of the samples in gnomAD overall v.3 (1,047/20,744). Moreover, classifying individuals based on self-reporting may not reflect the genetics basis, and force individuals to select “the best match” even though they wouldn’t identify as such.

### ADS calculation for a given dataset

The pipeline to calculate the ADS is available on GitHub, and we encourage anyone to use it. We strongly believe that it is essential to develop tools that can go to the data, rather than expecting the data to go to the tools, for example in a cloud environment. While we focus our research on the inclusion of systematically and structurally excluded populations, it is important to remember that in some cases, they are the owners of the data, and the data may not be deposited on public systems. Therefore, we encourage teams developing tools to make them easily accessible for use on local servers to ensure that they can be used widely, without discrimination.

### The ADS do not display population-specific information

Another limitation of displaying variant frequency per population is the lack of inclusion of populations for which sharing population information is not suitable. While the ADS does not inform about which population(s) carry the variant of interest, we believe it is up to each project, in partnership with the communities included in the dataset, to decide if they wish to display such information, and to whom. This is also a limitation of this technique. Indeed, with gnomAD or other databases separating their population, it is easy to determine if a variant is segregating in only one or a few population(s), and which ones. One solution is to associate, together with the ADS, the UMAP where each individual of the cohort is colored according to their genotypes for the variant of interest. However, before displaying such data, even if individuals cannot be identified, we strongly recommend engaging with the communities included in the analysis to ascertain their preference about such display.

### The ADS is dataset specific

The ADS is not a fixed value associated with a variant, it varies according to the data that were used to calculate the score. Indeed, a variant can be considered limited to one or a few populations (with a low ADS) when studying worldwide populations, but could be associated with a high ADS if only one of these populations is studied and the variant is spread across that population. For example, a variant present only in the Finish population, would have a low ADS if the UMAP is created with individuals coming from worldwide populations, but could have a high ADS if the studied population is only constituted of Finish individuals. Therefore, as for the allele frequency, it is important to indicate the scope of the dataset used to generate the ADS.

### Future directions

The generation of the ADS actively considers the issue of inclusivity among genomic datasets. To use this score in a clinical setting, for variants with a low ADS (i.e. a variant shared by individuals that are close genetically), it will be helpful to know if the patient would be from the same population as individuals carrying the variants, or be an outsider to those individuals. This could easily be achieved by sharing informative markers proximal to the variant location, but it requires thoughtful discussions to ensure that populations agree with sharing such information, or to develop a model that does not require sharing of informative markers. Following that implementation, it will be relevant to include the ADS into interpretation pipelines as genomic analysis leads the way toward personalized medicine.

Finally, as more projects are led by or work closely with systematically and structurally excluded communities, it is necessary to listen to them and their potential concerns about genomics derived information. For example, in this case, separating the individuals into populations and sharing population specific information was identified as a potential issue by some communities, and therefore, a method addressing their concerns was developed. More challenges may appear in future discussions and finding solutions to ensure that data ownership is conserved and that participants are at ease with the displayed information will be essential.

## Materials and methods

### Data

To develop and test the method, the final (phase 3) 2015 release of the International Genome Sample Resource (IGSR, formerly the 1000 Genomes Project), including 2,548 unrelated samples from 26 populations, representing five continental regions of the world, was used (Fairley et al., 2020). The IGSR dataset is available as multiple vcf.gz files (one per chromosome). Of note, the population labeling associated with each sample by the IGSR is based on an individual’s self-identification (1000 Genomes Project Consortium et al., 2015). The 2021 release of the IGSR, including 3,202 samples, was considered for use but ultimately excluded as some of the samples are related.

### Data preparation for UMAP generation

Due to the large number of variants and individuals, the release is split by chromosome. In order to generate the Uniform Manifold Approximation and Projection (UMAP) using data from all the chromosomes, 5% of the variants from each chromosome were randomly selected (--thin 0.05) and merged (--merge-list) into one file using PLINK (Chang et al., 2015). Then, using PLINK, the 15 first Principal Components (PCs) were calculated (--pca). Based on the 15 first PCs obtained by PLINK, the UMAP package from R was used to create a UMAP (McInnes et al., 2020). Details of the parameters used are available on the GitHub repository. The ggplot2 package was used to visualize the UMAP, and color the samples according to the associated population labels (Wickham, 2016).

### ADS calculation

Once the UMAP is generated using a subset of randomly selected data, the ADS is calculated for each variant with an minor allele count (AC) greater than or equal to two (--min-ac 2), as a variant heterozygous in only one individual (AC = 1) has an ADS of zero. After filtering the IGSR dataset, a total of 45,183,262 variants remained.

The ADS is calculated using a newly developed R script. First, the distance between each of the 2,548 individuals projected on the UMAP is included within a large matrix (2,548 columns x 2,548 rows). Then for each variant, the matrix is split into three matrices depending on the individual genotypes as depicted in S1 Figure (1. One matrix with the distances between individuals that are homozygous for the alternate allele, 2. One matrix with the distances between the individuals that are heterozygous and; 3. One matrix with the distances between individuals that are homozygous for the alternate allele and individuals that are heterozygous). The non-normalized ADS score is calculated based on individuals carrying at least one copy of the minor allele using the formula:

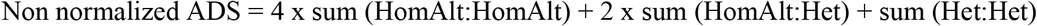

Finally, in order to normalize the ADS, the minimal and the maximal theoretical distances with the same number of individuals for each genotype category are calculated. These values are used to generate the normalized ADS, which can vary from 0 to 1.

### Comparison of the ADS with variant annotation and label score

#### Variant annotation

The variants were annotated using ANNOVAR following the user guide and using the default parameters (hg38) (Wang et al., 2010).

#### Label score

In order to compare the ADS with the current method consisting of reporting the allele frequency in each population, we calculated the label score for each variant. The label score is, for each variant, the number of populations with at least one individual carrying the alternate allele.

To assess if the variables had a distribution significantly different from a normal distribution, the Shapiro-Wilk normality test was used. As we can assume normality for all the variables (see full results in S6 Figure), we then used Pearson’s correlation.

### Analysis of a specific variant

To analyze the ADS of a given variant compared with ADS of variants with a similar allele frequency, the code uses PLINK (Purcell et al., 2007) and R. It loads a file with several variant IDs and creates the two outputs : The UMAP with individuals colored according to their genotype for the variant of interest and the relative position of the variant ADS within the ADS distribution for variants with a comparable allele frequency (+/− 10%).

### Data and code availability

To generate the ADS for a given dataset, the pipeline is available at : https://github.com/wassermanlab/ADS

It is composed of a shell script (1_ADS.sh, including the PLINK steps) and an R script (R_1_ADS.R, including the UMAP generation and the ADS calculation).

The ADS pipeline can be used with vcf, vcf.gz or ped/map files as input files.

To recreate the manuscript results, including the ADS calculation for the IGSR dataset and the figures, the code used is available on GitHub : https://github.com/wassermanlab/ADS/tree/master/IGSR_dataset_Manuscript/code

The dataset used was obtained from the IGSR website : http://ftp.1000genomes.ebi.ac.uk/vol1/ftp/data_collections/1000_genomes_project/release/20190312_biallelic_SNV_and_INDEL/

## Acknowledgments

We thank the Silent Genomes Project team and all the community members that are involved in this project for the interesting discussions that led to the development of this score. We also gratefully acknowledge past and present members of the Wasserman lab for helpful discussion. This work was supported by funding from the Canadian Institutes of Health Research (GP1-155868), Genome Canada and Genome British Columbia (275SIL), the Michael Smith Foundation for Health Research (17746) and the BC Children’s Hospital Foundation & Research Institute (KRZ48027).

