## Supplementary data for "Allele Dispersion Score: Quantifying the range of allele frequencies across populations, based on UMAP"

Cohort with 6 individuals : A, B, C, D E and F

Generation of a UMAP (Constant)

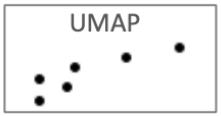


Matrix of distance between the 6 individuals (Constant)

| | A | B | C | D | E | F |
|---|---|---|---|---|---|---|
| A | x | x | x | x | x | x |
| B | dAB | x | x | x | x | x |
| C | dAC | dBC | x | x | x | x |
| D | dAD | dBD | dCD | x | x | x |
| E | dAE | dbE | dCE | dDE | x | x |
| F | dAF | dBF | dCF | dDF | dEF | x |

Individual's genotypes for varaint x

| Ind. | Genotype | Short |
|---|---|---|
| E, F | Homozygous reference | HomRef |
| C, D | Heterozygous | Het |
| A, B | Homozygous alternate | HomAlt |

Matrices for variant x

| | | Hom Alt | |
|---|---|---|---|
| | | A | B |
| HomAlt | A | x | x |
| | B | dAB | x |

sum (HomAlt:HomAlt) = dAB

| | | HomAlt | |
|---|---|---|---|
| | | A | B |
| Het | C | dAC | dBC |
| | D | dAD | dBD |

sum (HomAlt:Het) = dAC + dBC + dAD + dBD

| | | Het | |
|---|---|---|---|
| | | C | D |
| Het | C | x | x |
| | D | dCD | x |

sum (Het:Het) = dCD

Non-normalized ADS

Non normalized ADS = 4 x sum (HomAlt:HomAlt) + 2 x sum (HomAlt:Het) + sum (Het:Het)

**S1 Figure : Overview of the method used to calculate the non-normalized ADS.**

Calculation of the non-normalized ADS for variant ‘x’ in a population of six individuals : A, B, C, D, E and F.

First, a UMAP is generated with all individuals, then a matrix including the distance between each of the 6 individuals projected on the UMAP is created. The UMAP and the distance matrix are constant for the calculation of the ADS of each variant. Then for each variant, the matrix is split into three matrices depending on each of the individual genotypes. In this example, individuals A and B are homozygous for the alternate allele, individuals C and D are heterozygous and individuals E and F are homozygous for the reference allele. The three matrices are: 1. One matrix with the distances between individuals that are homozygous for the alternate allele (individuals A and B). 2. One matrix with the distances between the individuals that are heterozygous (individuals C and D). 3. One matrix with the distances between individuals that are homozygous for the alternate allele (individuals A and B) and individuals that are heterozygous (individuals C and D). The non-normalized ADS score is calculated based on individuals carrying at least one copy of the minor allele using the formula presented at the bottom. After this step, a normalization is performed as explained in the main manuscript.

**A.**
**(Figure 2B)**

| Minor Allele Frequency (MAF) bins | Number of variant |
|---|---|
| [0,0.01] | 31,795,321 |
| (0.01,0.02] | 2,899,058 |
| (0.02,0.03] | 1,369,198 |
| (0.03,0.04] | 849,808 |
| (0.04,0.05] | 611,100 |
| (0.05,0.06] | 475,106 |
| (0.06,0.07] | 394,132 |
| (0.07,0.08] | 342,104 |
| (0.08,0.09] | 309,531 |
| (0.09,0.1] | 280,330 |
| (0.1,0.11] | 261,468 |
| (0.11,0.12] | 242,084 |
| (0.12,0.13] | 231,004 |
| (0.13,0.14] | 216,293 |
| (0.14,0.15] | 206,049 |
| (0.15,0.16] | 194,561 |
| (0.16,0.17] | 188,003 |
| (0.17,0.18] | 183,575 |
| (0.18,0.19] | 171,071 |
| (0.19,0.2] | 169,974 |
| (0.2,0.21] | 166,303 |
| (0.21,0.22] | 159,770 |
| (0.22,0.23] | 153,022 |
| (0.23,0.24] | 149,601 |
| (0.24,0.25] | 145,685 |
| (0.25,0.26] | 139,566 |
| (0.26,0.27] | 138,049 |
| (0.27,0.28] | 135,096 |
| (0.28,0.29] | 135,004 |
| (0.29,0.3] | 131,117 |
| (0.3,0.31] | 130,282 |
| (0.31,0.32] | 128,276 |
| (0.32,0.33] | 125,773 |
| (0.33,0.34] | 123,882 |
| (0.34,0.35] | 120,683 |
| (0.35,0.36] | 120,781 |
| (0.36,0.37] | 118,947 |
| (0.37,0.38] | 117,521 |
| (0.38,0.39] | 115,858 |
| (0.39,0.4] | 116,930 |
| (0.4,0.41] | 114,536 |
| (0.41,0.42] | 115,235 |
| (0.42,0.43] | 112,144 |
| (0.43,0.44] | 112,023 |
| (0.44,0.45] | 111,693 |
| (0.45,0.46] | 112,979 |
| (0.46,0.47] | 113,986 |
| (0.47,0.48] | 110,721 |
| (0.48,0.49] | 109,767 |
| (0.49,0.5] | 108,262 |
| Total | 45,183,262 |

**B.**
**(Figure 2C)**

| Minor Allele Frequency (MAF) bins | Number of variant |
|---|---|
| [0,0.0005] | 8,289,572 |
| (0.0005,0.0015] | 11,498,871 |
| (0.0015,0.0025] | 3,875,941 |
| (0.0025,0.0035] | 2,263,416 |
| (0.0035,0.0045] | 1,555,669 |
| (0.0045,0.0055] | 1,369,976 |
| (0.0055,0.0065] | 884,350 |
| (0.0065,0.0075] | 729,693 |
| (0.0075,0.0085] | 611,307 |
| (0.0085,0.0095] | 527,265 |
| (0.0095,0.0105] | 457,672 |
| (0.0105,0.0115] | 404,278 |
| (0.0115,0.0125] | 361,957 |
| (0.0125,0.0135] | 324,661 |
| (0.0135,0.0145] | 293,720 |
| (0.0145,0.0155] | 267,330 |
| (0.0155,0.0165] | 292,795 |
| (0.0165,0.0175] | 221,491 |
| (0.0175,0.0185] | 202,727 |
| (0.0185,0.0195] | 190,497 |
| (0.0195,0.0205] | 176,127 |
| (0.0205,0.0215] | 163,925 |
| (0.0215,0.0225] | 157,367 |
| (0.0225,0.0235] | 149,060 |
| (0.0235,0.0245] | 140,159 |
| (0.0245,0.0255] | 130,846 |
| (0.0255,0.0265] | 148,125 |
| (0.0265,0.0275] | 115,074 |
| (0.0275,0.0285] | 109,831 |
| (0.0285,0.0295] | 107,763 |
| (0.0295,0.0305] | 103,583 |
| (0.0305,0.0315] | 96,850 |
| (0.0315,0.0325] | 92,798 |
| (0.0325,0.0335] | 89,100 |
| (0.0335,0.0345] | 85,532 |
| (0.0345,0.0355] | 82,020 |
| (0.0355,0.0365] | 94,466 |
| (0.0365,0.0375] | 75,037 |
| (0.0375,0.0385] | 74,444 |
| (0.0385,0.0395] | 70,506 |
| (0.0395,0.0405] | 68,420 |
| (0.0405,0.0415] | 67,377 |
| (0.0415,0.0425] | 65,411 |
| (0.0425,0.0435] | 63,893 |
| (0.0435,0.0445] | 61,335 |
| (0.0445,0.0455] | 58,881 |
| (0.0455,0.0465] | 57,367 |
| (0.0465,0.0475] | 66,026 |
| (0.0475,0.0485] | 54,154 |
| (0.0485,0.0495] | 54,429 |
| (0.0495,0.05] | 21,391 |
| Total | 37,524,485 |

**S2 Table : Number of variants per bin used for figure 2B and 2C of the main manuscript.**
Figure 2B and 2C represents the ADS distribution according to the Minor Allele Frequency (MAF). In Figure 2B, all variants (0 < MAF < 0.5) are binned according to their MAF in bins of 0.01, and the number of variants per bin is represented here in supplementary figure 2A. In Figure 2C, variants with a MAF < 0.05 (The same variants as the ones present in the first 5 bins of supplementary table 2A) are binned according to their MAF in bins of 0.01, and the number of variants per bin is represented here in supplementary figure 2B.

| Variants annotation | Number of variants |
|---|---|
| ncRNA_exonic;splicing | 53 |
| exonic;splicing | 126 |
| UTR5;UTR3 | 317 |
| ncRNA_splicing | 1,014 |
| splicing | 2,332 |
| upstream;downstream | 12,402 |
| UTR5 | 90,622 |
| ncRNA_exonic | 175,396 |
| upstream | 283,505 |
| downstream | 311,895 |
| exonic | 412,124 |
| UTR3 | 508,899 |
| ncRNA_intronic | 2,811,900 |
| intronic | 17,215,489 |
| intergenic | 23,357,104 |
| total | 45,183,178 |

**S3 Table : Number of variants per annotation used for Figure 2D of the manuscript.**
Number of variants per annotation class as plotted in manuscript figure 2D, which represents the ADS distribution according to variants annotation.

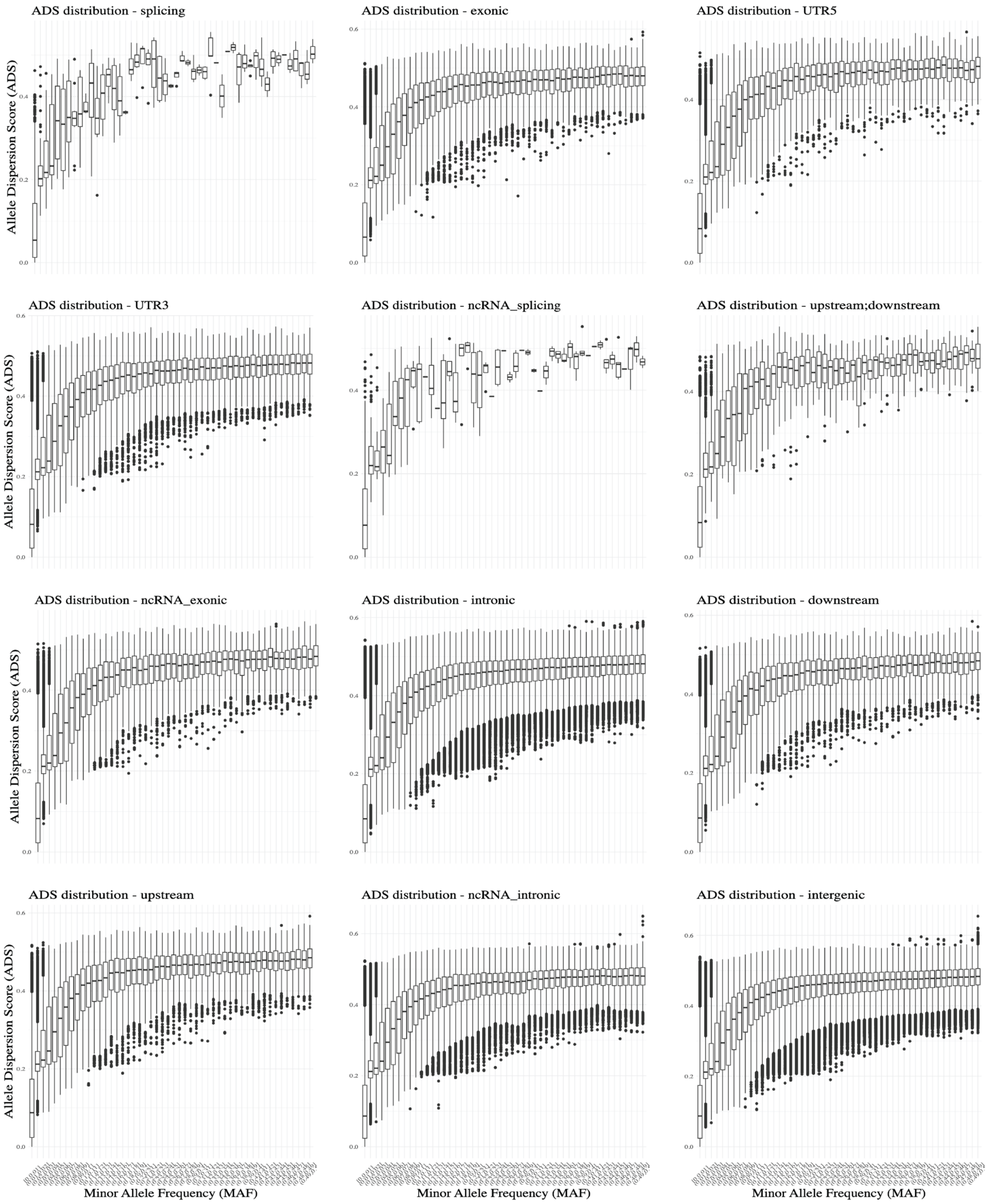


**S4 Figure : ADS distribution per MAF for each annotation class.**
For each panel, the variants were separated according to their annotation class (Annotation performed with ANNOVAR). Only annotations with more than 500 variants are represented.
For each variant class, the boxplot represents the ADS distribution according to the Minor Allele Frequency (MAF). Variants are binned according to their MAF (bins of 0.01).
The pattern of ADS distribution compared to the MAF appears to be the same across all variant classes.

A.

| Continental label | Number of individuals |
|---|---|
| AMR | 348 |
| SAS | 492 |
| EAS | 515 |
| EUR | 522 |
| AFR | 671 |
| Total | 2,548 |

B.

| Population label | Number of individuals |
|---|---|
| ASW | 61 |
| MXL | 64 |
| PEL | 85 |
| BEB | 86 |
| MSL | 90 |
| CLM | 95 |
| PJL | 96 |
| ACB | 97 |
| CEU | 99 |
| KHV | 99 |
| CDX | 100 |
| ESN | 100 |
| GBR | 100 |
| ITU | 102 |
| STU | 102 |
| LWK | 103 |
| PUR | 104 |
| CHS | 105 |
| FIN | 105 |
| JPT | 105 |
| CHB | 106 |
| GIH | 106 |
| IBS | 107 |
| YRI | 107 |
| TSI | 111 |
| GWD | 113 |
| Total | 2,548 |

**S5 Table : Number of individuals per continental (A) and population (B) label in the IGSR dataset.** Within the IGSR data, each individual is associated with two population labels: i) a continental label or super-population (n = 5); and ii) a population label (n = 26). The number of individuals per population is not evenly distributed. Of note, the population labeling associated with each sample by the IGSR is based on an individual's self-identification (1000 Genomes Project Consortium et al., 2015).

Shapiro-Wilk normality test

| Variable | W | p-value | Conclusion |
|---|---|---|---|
| Continental label score | 0.9868 | 0.9672 | Not significantly different from normal distribution |
| Population Label score | 0.9583 | 0.3592 | |
| Mean ADS | 0.9838 | 0.9432 | |

Pearson's product-moment correlation

| Figure | Variable 1 | Variable 2 | df | r coefficient | p-value | Conclusion |
|---|---|---|---|---|---|---|
| 3A | Continental label score | Mean ADS | 3 | 0.968769 | 0.006594 | Positive correlation and significant relation |
| 3B | Population Label score | Mean ADS | 24 | 0.9806424 | < 2.2e-16 | Positive correlation and significant relation |

**S6 Figure. Details of the statistical results obtained for the comparison of the ADS and the label score.** The Shapiro-Wilk Normality test indicates a variable distribution that differs significantly from a normal distribution. As the three variables are not significantly different from normal distribution, a Pearson's product-moment correlation was applicable to study the correlation of the ADS and each label score. The ADS correlates positively with each of the label scores and the relationships are significant.
